# Bacterial Growth Stimulation and Antifungal Effects Of The Essential-oil-less-extracts Of The Food Spice *Dysphania ambrosioides*

**DOI:** 10.1101/2020.02.04.934885

**Authors:** Lucía Nitsch-Velásquez

## Abstract

1

*D. ambrosioides* leaves (DaL) are utilized as food spice. DaL infusion is utilized as antihelminthic in traditional medicine, the extracted essential oil (EO) has been repurposed as biopesticide, *in vitro* activities include nematocidal and phytotoxic activities by hydrophillic compounds from leaves and the roots. As Da might be a good candidate for circular economy, more potential applications of the essential–oil–less–Da extracts were pursued by applying green chemistry based extraction methods.

DaL extracts were prepared by the autoclave method for the sterile–essential–oil–less aqueous extract (SALAEL–Da), butanol fractionating for saponins extraction (SAP) and ethanol boiling method for the saponin–free extract (EtOH–Da). Their effects over clinical isolates of fungi (*Candida albicans* / CA), gram–negative bacteria (*Erwinia carotovora* / ErC) and –positive (Methicillin resistant *Staphylococcus aureus* USA 300 / MRSA–USA–300). were explored.

**The raw extracts stimulated the bacterial growth of all strains in the pre-screening phase.** SALAEL-DaL (at 25 mg/mL) estimulated ErC growth, reducint its doubling time by 35%, microdilutions of EtOH-DaL (at 180 mg/mL) stimulated the growth of MRSA–USA–300 even in the GEN presence at sub-lethal concentrations (MIC_*GEN*_=1.75*µ*g/mL). SALAEL-Da (at 137 mg/mL) inhibited the growth of CA in agar dilutions, and its fraction SAP showed a moderated fungistatic effect at 100 mg/mL in disk diffusion pre-screening tests. SAP fraction may partially account for the observed antifungal activity. The essential–oil–less–Da aqueous extracts analyzed contained bacterial growth stimulating and antifungal components. Further investigation may lead to commercial opportunities for probiotics and antifungals.

**Importance:** The leaves of *D. ambrosioides* (Da) are utilized as food spice and its infusion as antihelminthic in Latin American folk medicine, with a wide variety of *in vitro* bioactivities reported. The most studied is Da essential oil (Da–EO) and has been repurposed as insect repellent and pesticide. Which would leave the extracted plant material as a potential raw material for other products. Additionally, Da could be an interesting candidate tobe produced at high–scale in rural communities. The ecofriendly extraction processes yielded an aqueous extract (free of the EO) that enhances the bacterial growth rate of two bacterial strains, which can eventually be useful in the probiotics industry. This extract also inhibited the growth of the opportunistic fungi *Candida albicans*, becoming a potential source of new antifungals. Further investigation may lead to a circular economy of the agroindustry around *D. ambrosioides*, probiotics and antifungals, among others.

## 3 Introduction

*D. ambrosioides* leaves are utilized in folk medicine around the globe—*e.g.*, Navajo, Mayan, Peruvian, Indian, Chinese, Moroccan, Egyptian cultures—and as food spice in Latin America^1–7^, *e.g.*, Da leaves are added to beans to prevent flatulence^8^. Da is an ecological opportunist plant, endemic of Central America, but has been so successfully introduced worldwide^9^ that in some countries it is considered a weed.^**?**^ a is usually co–cultivated in family gardens in rural communities and sold in local markets for traditional utilization, The EO has being repurposed from the initial antihelmintic and antiprotozoal ‘Baltimore oil’^5,10–12^ to applications as biopesticide^13–19^ and weed control^**?**^. A wide variety bioactivities of Da extracts have been screened^7,20,20**?**–22^. The aereal parts have shown antibacterial^23–25^ and antifungal^22^ activities, genotoxicity^12^ and slight hepatotoxicity^26^ (in all the studies the extracts still contained the EO). The EO antimicrobial activities (ascaridole is its main component)^27^. MacDonald et al.^5^ found that the ascaridole–free–aqueous extract is also nematocidal (*e.g.*, bioactive hydrophyllic compounds), suggesting that the traditional utilization of *D. ambrosioides* infusion as vermifuge appears to be safer than the purified EO. *D. ambrosioides* extracts are rich in secondary metabolite families (glucosides, flavonoids, triterpenes, saponins, alkaloids probably from the piperidine family, among others)^7,23,28^ with structural elucidation studies pending for most of them.

### The whole Da plant could be exploided under a circular economy^29^ framework

It can be envisioned as follows: first, the essential oil is extracted for perfumery or biopesticide industries, then the produced waste (the extracted plant material) becomes the raw material for other products (*e.g.*, source of nematocidal agents, antimicrobials, food additives, nutraceuticals, biopesticides), and the leftover can become another type of food or organic fertilizer^30^. Additionally, due to its eco–adaptability and the potential whole plant explotation, Da could be an interesting candidate for livelihood diversification in remote communities^28,31–34^. In order to pursue the expansion of Da exploitation, it is needed to determine other properties of its essential oil–less–extracts obtained by green chemistry extraction processes. Their potential antimicrobial activities against representative gram negative, gram positive bacteria, and fungi (*Erwinia carotovora*—ErC, Methicillin Resistant *Staphylococcus aureus* USA–300—MRSA–USA–300 and *Candida albicans*—CA, respectively) were targeted in this study.

## 4 Results

All the polar essential oil–less–extracts of *D. ambrosioides* stimulated the growth of ErC and MRSA–USA–300 in the pre-screening phase. In the FICI–pre–screening, the ethanolic extract (EtOH-DaL) stimulated the MRSA–USA–300 growth in the presence of sublethal gentamicin concentrations (MIC_*GEN*_=1/75 *µ*g/mL, see Table 1.a.) but did not affect the MIC_*GEN*_ *per se*. The ErC growth curve experiments with the sterile–aqueous extracts of *D. ambrosioides* leaves (SALAEL–Da) also showed the bacterial growth stimulation effect. In Table 1.b are tabulated the estimated doubling times (*t*_*D*_) of ErC for different SALAEL–Da concentrations, notice that *t*_*D*_ was reduced by a 35% at the extract concentration of 25 mg/mL, with differentiated and non linear response to concentration, for instance at 100 mg/mL *t*_*D*_ was decelerated.

**Table 1:**
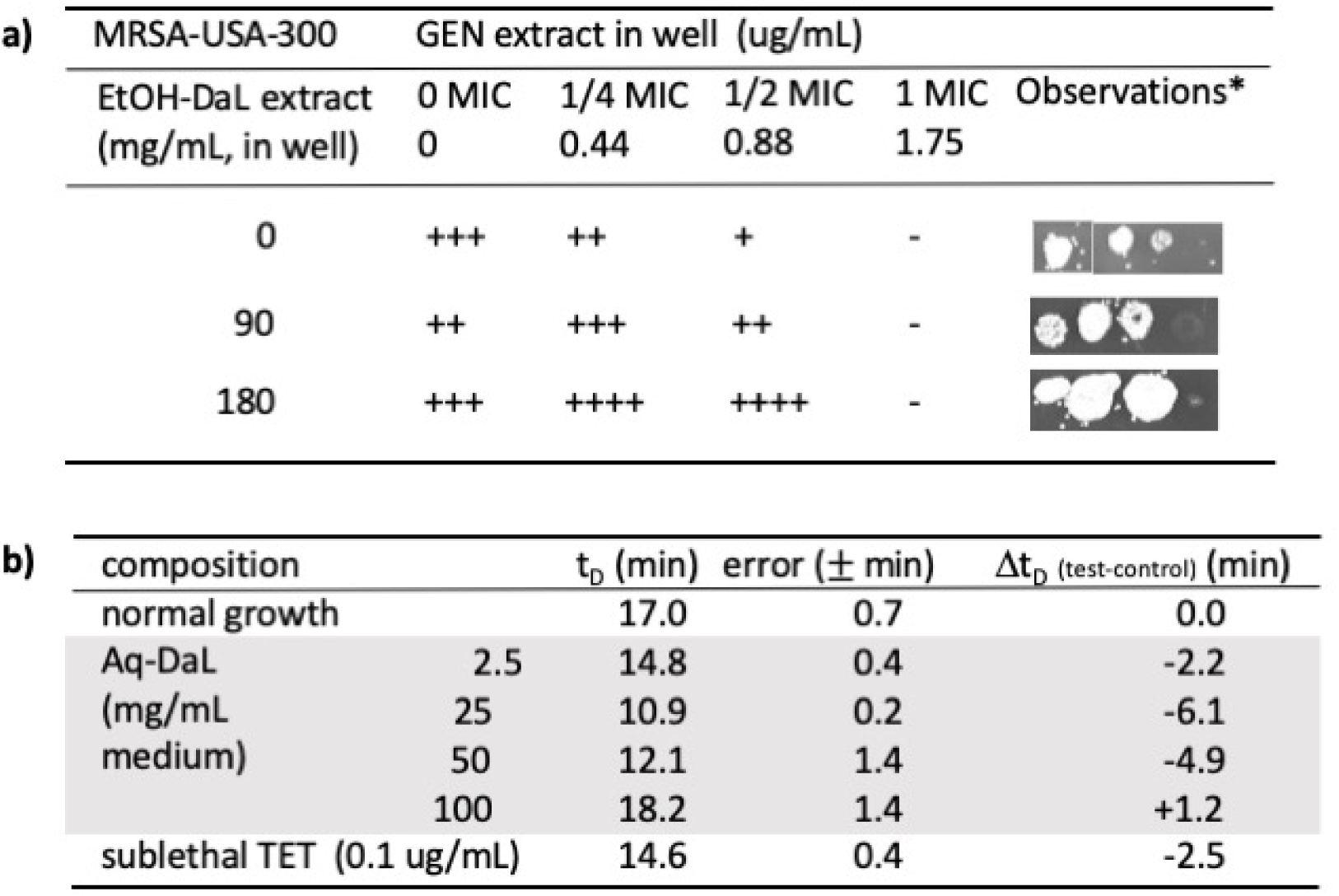
Effects of *D. ambrosioides* extracts over gram positive and negative bacteria. **a)** Exploratory fractional inhibitory concentration index of EtOH-DaL and gentamicin, observe the overgrowth of tests compared to the normal growth control at sublethal GEN concentrations. **b)** Effects of SALAEL-Da extracts over the dobling time of *E. carotovora* determined by the growth curve and Gomptertz model adjustment. Notice the reduction exerted by the extract at 25 mg/mL. Both extracts yielded 2% (w/w) each from the dried plant material.

| MRSA-USA-300 | GEN extract in well (ug/mL) |  |  |  | Observations* |
| --- | --- | --- | --- | --- | --- |
| EtOH-DaL extract (mg/mL, in well) | 0 MIC | 1/4 MIC | 1/2 MIC | 1 MIC |  |
|  | 0 | 0.44 | 0.88 | 1.75 |  |
| 0 | +++ | ++ | + | - |  |
| 90 | ++ | +++ | ++ | - |  |
| 180 | +++ | ++++ | ++++ | - |  |

**b)**
| composition | | $t_D$ (min) | error (± min) | $\Delta t_D$ (test-control) (min) |
| --- | --- | --- | --- | --- |
| normal growth |  | 17.0 | 0.7 | 0.0 |
| Aq-DaL | 2.5 | 14.8 | 0.4 | -2.2 |
| (mg/mL | 25 | 10.9 | 0.2 | -6.1 |
| medium) | 50 | 12.1 | 1.4 | -4.9 |
|  | 100 | 18.2 | 1.4 | +1.2 |
| sublethal TET (0.1 ug/mL) |  | 14.6 | 0.4 | -2.5 |

**The SALAEL–Da inhibited the growth of *Candida albicans* clinical isolate (CA)** at the concentration of 135 mg/mL agar, see Table 2. An alkaloidal saponin fraction (SAP) was detected by positive Draggendorf’s, hemolysis and foam tests, being extractable with butanol. The estimated hemolysis index for SAP was of 22,500, while showed moderated fungistatic activity in the pre–screning phase (∼0.1 mm growth inhibition halum, see Fig. 1.a & b).

**Table 2:**
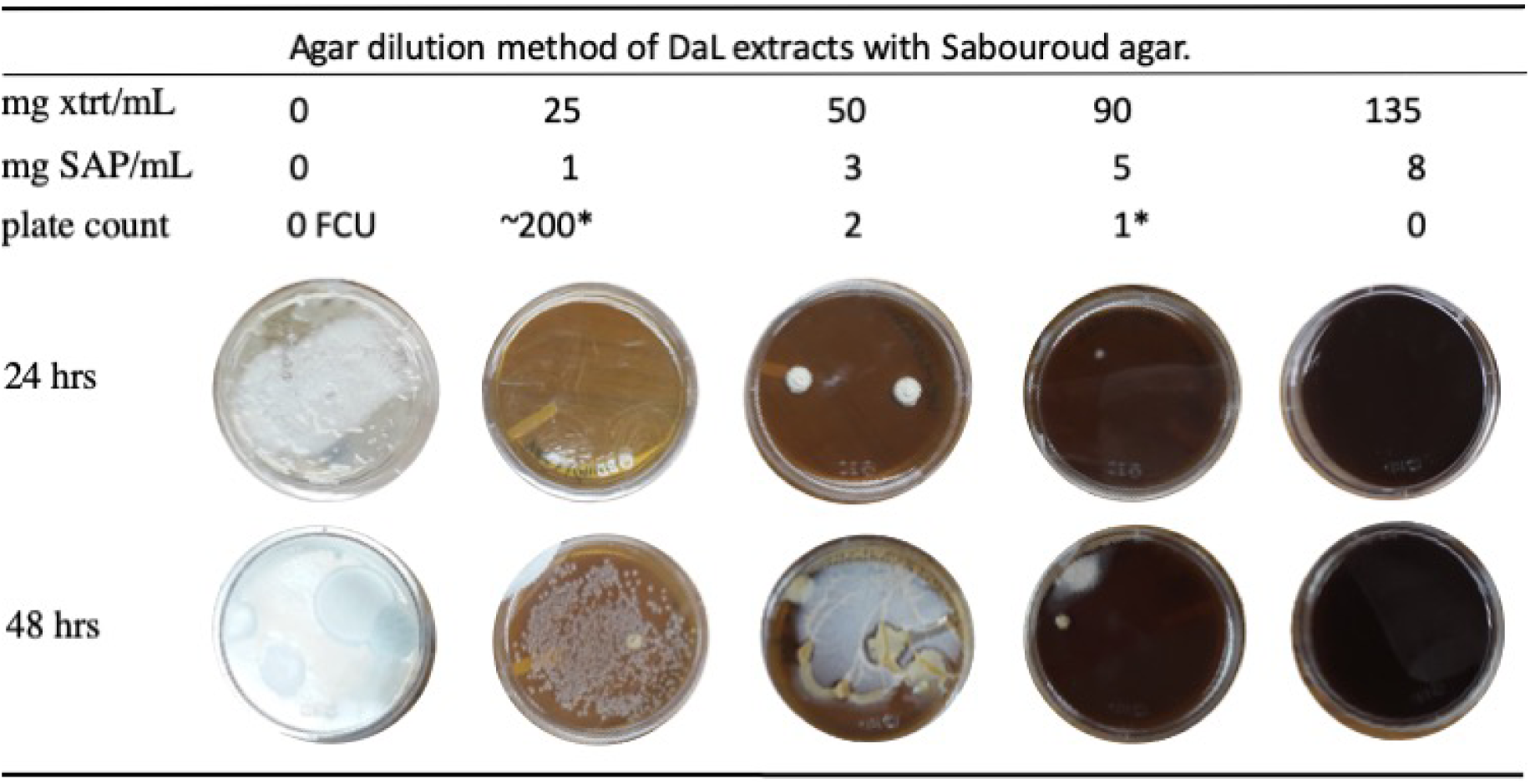
Inhibitory effect of the sterile essential oil–less aqueous extract from *D. ambrosioides* leaves (SALAEL–Da) over the growth of the opportunistic yeast *Candida albicans* (clinical isolate, CA–ci). Inhibition occurs at the concentration of 135mg/mL SALAEL–Da. Notice the faster development of selected former colony units in the growth medium with 25 mg/mL SALAEL–Da. Saponin presence in the extract was detected by the haemolysis test. SAP yield was 3% (w/w) from the raw extract.

| Agar dilution method of DaL extracts with Sabouroud agar. |  |  |  |  |  |
| --- | --- | --- | --- | --- | --- |
| mg xtrt/mL | 0 | 25 | 50 | 90 | 135 |
| mg SAP/mL | 0 | 1 | 3 | 5 | 8 |
| plate count | 0 FCU | ~200* | 2 | 1* | 0 |
| 24 hrs |  |  |  |  |  |
| 48 hrs |  |  |  |  |  |

**Figure 1:**
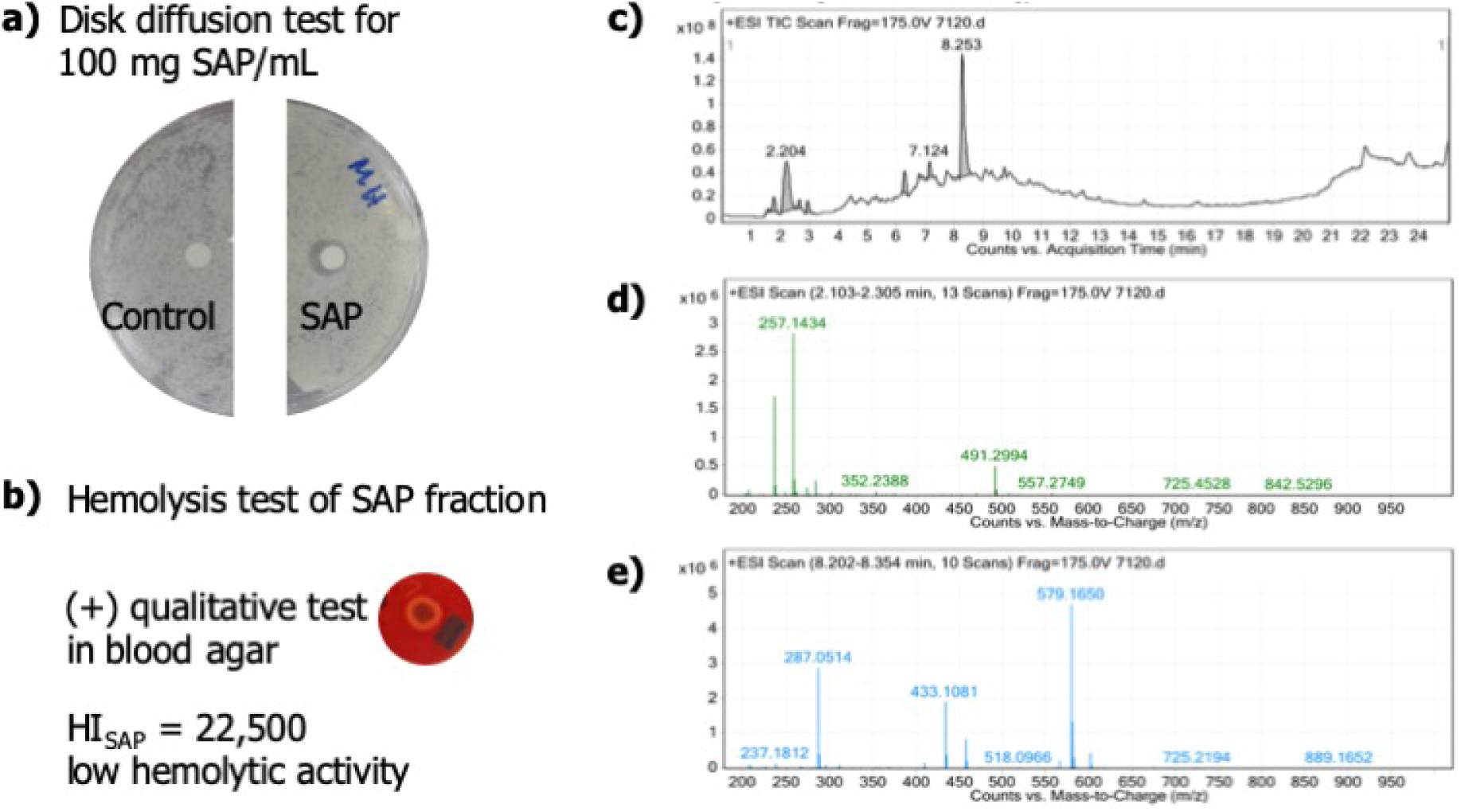
Saponin fraction (SAP) general characterization: **a)** Moderated fungistatic activity against CA in Mueller-Hinton agar. **b)** Qualitative and quantitative hemolysis tests. **c)** LC–MS chromatogram of SAP (10*µ*g/mL). **d)** ESI+ full scan of t_*R*_ 2.103-2.305 min. **e)** ESI+ full scan of t_*R*_ 8.202-8.354 min.

The saponin fraction yielded the LC-MS is shown in Fig. 1.c to e, the two MS–full scan included show the fraction of two chromatographic peaks are included. The mass-to-molecular formula analysis showed that among its components is at least one nitrogen, for instance *m/z* 257.1434 can be related to the molecular formula C_5_H_19_N_7_O_5_, +3.2 ppm, other potential candidates also contained nitrogen (in agreement with the positive alkaloids test).

## 5 Discussion

The extracts were referred as essential–oil–less extracts of *D. ambrosioides* due to the extraction processes, by which is expected the original EOs would be removed by heat. Even though the processes may yield reaction products (*e.g.*, hot hydrolysis could release free volatile carboxylic acids from their ester forms), autoclave conditions as part of the selected extraction method are ecofriendly, yields sterile products and can potentially be applied at industrial level. The bioactive concentrations of all the extracts were in the concentration range of extracts with medium–to–high potential for containing bioactive components for which further purification is recommendable^35–38^.

The EtOH–DaL and SALAEL–Da stimulated the growth of MRSA–USA–300 and ErC respectively, but the linearity response should be investigated (see Table 1). The fact that promotes the bacterial growth of gram–positive and –negative makes *D. ambrosioides* a potentially attractive material for the probiotics ^**A**^ and biotechnology industries^42–45^, *e.g.*, probiotics based on *Lactobacillus* sp., a gastrointestinal tract flora component^46^, ErC strains EC UV4 for 2,3–butanediol production from corncob^47^, and *Enterobacter cloacae* IIT–BT08 for biohydrogen production^48^.

**The CA growth inhibition of the SALAEL-Da might be partially** explained by the alkaloidal saponin(s) presence, but other—unknown—components contributed to its bioactivity^49^. The SAP relatively low hemolysis index is a favorable feature for drug development, for instance as potential vaccine adjuvant^50^. It should be mentioned the unexpected physiological estimulation in the agar dilution method is shown Table 2, 25 mg/mL SALAEL–Da. Usually after 24 hrs, CA would be as a yeast form, but for this case the observed physiological state would be normally observed after 48 hrs or more. Their effect on bacterial strains should be explored, even though saponins have not been reported as bacterial growth stimulants, the potential utilization of saponins as adjuvants of antifungal agents would require this verification step, among others (*e.g.*, citotoxicity).

**Potential future research directions regarding SALAE–Da extracts** and its derivatives as potential microbiology commercial products are: a) extract’s composition and structural elucidation of isolable compounds, b) antifungal activity, c) effects on fungi development, d) potential antimicrobial adjuvant of the saponin fraction, e) stimulation of the bacterial growth rate of both the raw extracts and their fractions in bacteria of industrial interest.

**As the probiotics’ market is on the** grow^**B** 51^ requiring the industrial processes optimization, *e.g.*, by adding bacterial elicitors to the growth medium in order to speed up the production. Also, the demand of ‘smart’ probiotic products —such as cosmetics and skin medications— may be a commercial trend that might be becoming a need for the public health^40^. The repurposing of *D. ambrosioides* might be part of that emerging market in the format of a circular and socially responsible economy.

## 6 Materials and Methods

### Materials

*Antibacterial agents*: gentamicin (GEN, Sigma, 10mg/mL commercial sterile solution), tetracycline (TET, Merck, 99,9%, solid); GEN *Ez strips*^*R*^ for minimal inhibitory concentration determination (BioMerieux). *Surfactant agents*: dodecyl sodium sulfate (SDS, anionic surfactant, Sigma, 98%, solid), alconox (anionic detergent, Sigma–Alconox Inc.), saponin (non–ionic surfactant, plant derived, Sigma, 8–25%). *Solvents*: ultrapurified water (mqH_2_O, MilliQ system, resistivity of 18.2 Mohms*cm), distilled water (dH_2_O, Salvavidas), Ethanol 95% (Merck), methanol 98% (Merck), dimethyl sulfoxide (DMSO, Merck, 99.9%), phosphate buffered solution standard tablets (PBS, Sigma: 0.01 M phosphate buffer, 0.0027 M potassium chloride and 0.137 M sodium chloride, pH 7.4, at 25 °C). *Growth mediums* Mueller-Hinton agar and broth (Merck), brain heart infusion agar and broth (Hardy), Sabouroud agar (Merck), blood agar (Merck) were prepared according to the supplier recommendations. *Strains*: Methicillin Resistant *Sthapylococcus aureus*–USA–300 (MRSA–USA–300), *Enterobacter cloacae*–TET susceptible, environmental isolate; *Erwinia carotovora*–TET susceptible, plant isolate; *Candida albicans*, clinical isolate.

### Extracts

*Plant collection and identification*: *D. ambrosioides* was cultivated in the NGO garden, collected and botanically verified (Voucher 005), shadow dried, and leaves were separated. Note: the plant was associated to a rust type fungi. *Ethanolic extract* (EtOH-Da): About 1kg of plant material was extracted with one gallon of 95% ethanol (Laksochem), boiled for 45 min, cloth filtrated and hot evaporated under atmospheric pressure. *Sterile aqueous extracts* (SALAEL-Da): Each extraction batch consisted of about 1.5 kg in 3L distilled water distributed in 250 mL erlenmeyers. Standard autoclave conditions for liquid medium were applied. Extract was cloth filtrated and hot evaporated under atmospheric pressure, stored at 4°C.Notice that the sample was considered as an ascaridol-less extract or that the essential oil was removed from the sample during the high–pressure and high–temperature extraction process. *Saponin fraction* (SAP): 20 g of extracts were exhaustively extracted with butanol (Merck, 95%), monitored by hemolysis-TLC.

### Alkaloids qualitative test

10 *µ*L of the saponin fraction were tested with the Draggendorf’s solution^53^.

### Growth curve

*Mixtures preparation*: for extract related tests, the preparation of BHI broth was modified, such that the growth medium components were at the manufacturer’s recommended concentration, after adding the plant extract. 1mL of the seeding bacterial dilution at 0.5 McFarland was diluted to 100 mL final volume, with growth medium mixture (GMMx). The expected final UFC would be 1.5 * 10^6^ Former Colony Units / mL GMMx, being the bacterial’s physiological at the linear growth phase. *Sampling*: The bacterial growths were sampled in one hour intervals, under sterile conditions, taking two 100*µ*Laliquots from each mixture; they were directly transferred to the plastic cells assigned to each mixture, the same cell was utilized along the experiment. Concentration was determined at 600 nm.^54,55^.

### FICI pre-screening

The test solution was constituted of 90 *µ*Lof MH broth with mixtures of GEN (at 0, 1/4, 1/2 and 1 MIC as final well concentrations) with ethanolic extract (0, 100, 200 mg/mL, idem before) and were seeded with 10 *µ*Lof 10^6^ MRSA–USA–300 dispersion, then incubated 24 hrs in ELISA plates at 32°C. This was followed by the reincubation of 10 *µ*Lof test solution for additional 24 hrs. Samples were run in duplicate.

### Fungistatic activity against *Candida albicans*

*Agar dilution method:* The CLSI protocol for yeast MIC determination by the agar dilution method (ADM)^54,56^ was followed with several modification. Sabouroud agar was prepared according to the supplier recommendation, taking in to account the volume contribution from the dissolved extract as if it was water only. The controls included were: medium sterility (one plate) and normal growth (triplicate), tests were run in triplicate. *Disk diffusion method* ^57^: Disks of Whatman No. 60 filter paper, 0.5 mm diameter were utilized.

**Hemolysis qualitative tests** were performed in blood agar^58^ and thin layer chromatography^59^ methods (for the latter, the blood solution concentration was 10 fold higher). *Spetrophotometry quantification*^60,61^ Controls included were full hemolysis by 10% alconox solution (final concentration 0.1%), distilled H_2_O as hypotonic media. Testing extracts at final concentration range was 0.1–20 mg/mL. The haemolysis percentage^50^ and hemolysis index were determined (value of Saponin D concentration for 100% hydrolysis of 2 mL blood)^60^.

### Data analysis

The microbial concentration was transformed to biomass of Former Colony Units or FCU from OD_600_ ^62^. Descriptive statistic of the determinations of the minimal inhibitory concentrations were calculated with Excel v2016, Microsoft Windows software. Bacterial growth curves were analyzed applying the Gomptertz model for bacterial growth utilizing Mathematica software^48,62–65^. All raw data was included in all the analysis, plots of the normalized data were prepared by subtracting the absorbance due to the extract’s and growth medium components. The effect of a given treatment over the doubling time was classified as significantly different from the normal growth based on the simplified Pocock’s statistic test^66^.

### LC–MS analysis

Scripps laboratory commercial services were contracted. LC–MS TOF Agilent 6200 series B.05.0, ESI, positive mode, 175V, 20*µ*Lwere injected in a C18 2.1×150mm column, applying polarity gradient of water–0.1%formic acid and ACN–0.1% formic acid, from 95:5 to 5:95 in 20 min, with a 5 min hold at 5:95, total runtime of 25 min.

## 7 Acknowledgments

To the Association of Medical School Microbiology and Immunology Chairs (AMSMIC), Dr. Anthony Campagnari, Dr. Nicole Luke (UB) and to Universidad del Valle de Guatemala for sharing their strains and research facilities. Dr. Sara Barrios (PERA) for identification of specimes utilized in this study. Dr. Diana Aga and Dr. Troy Wood (UB), Dr. Giovanni Sindona, Dr. Domenico Taverna, Dr. Galileo Violini (Universitat della Calabria, Italy), and Victor Jimenez (UVG) for sharing their expertise, Dr. Dalia Lau, Dr. Krisztina Rios-Fulop (UVG) for helpful discussions, and Ma. Luisa Mendizabal, David Archila, Miguel Morales, Nina Figueroa, and Elena Ortiz–Osejo (UVG) for their technical assistance.

## Footnotes

A A standard definition of probiotics is not available yet, but a definition of probiotics is: *live microorganisms* ^*39,40*^, *that are administered in adequate amounts* ^*40,41*^, *with an intended therapeutic effect for the host* [(i.e.] *probiotics for clinical use* [in humans] ^39,40^.

B It is expected that the global market for probiotics will reach about USD 66 billion by 2024^51^. The probiotics’ market is triggering a set of *unique scientific, translational and regulatory challenges* ^52^.

